# Goal-directed shaping of cortical waves

**DOI:** 10.1101/2024.09.10.612344

**Authors:** Anton A. Dogadov, Daniel E. Shulz, Valérie Ego-Stengel, Isabelle Ferezou, Luc Estebanez

## Abstract

At the surface of the cerebral cortex, the dynamics of brain activity at the mesoscopic scale are characterized by waves of synchronized neuronal activity. These waves have been shown to impact the processing of sensory information, but can they be actively shaped by the subject in a goal directed manner? To address this question, we designed a fast widefield optical brain-machine interface for mice that can detect and reinforce individual traveling waves, which follow a specific displacement at the surface of the somatosensory cortex. Trained mice learned to generate these Conditioned Waves, which became progressively more stereotyped. The Conditioned Waves resulted from a reshaping of the cortical activity associated with limb movements, which included a sharp pre-movement cortical suppression that emerged with learning. Our work demonstrates that traveling cortical waves of neuronal activity can be subject to operant control. It provides evidence for the plasticity and functional relevance of large-scale cortical dynamics, and establishes a new paradigm for manipulating mesoscale brain activity.

## INTRODUCTION

During awake, active behavior, the activity of cortical neurons is typically described as dominated by asynchronous, fast fluctuations in membrane potential (Steriade et al. 2001; Poulet and Petersen 2008). However, the advent of high-resolution recording methods applied to large cortical territories has revealed that such “active” cortical state is not uniform but rather shows patterns of synchronous firing resulting in a constant flow of waves traveling across the cortical network (Ma et al. 2016; Mohajerani et al. 2010; Luczak et al. 2009; Shahsavarani et al. 2023). These waves tend to move from sources, often located in the primary sensory areas, towards sinks — some located in associative areas (Mohajerani et al. 2013; Liang et al. 2023). Time averaging of this mesoscale activity during trained behaviors revealed networks of sensory, motor and high-order cortical areas that are activated in a highly structured fashion as the behavior unrolls (Allen et al. 2017; Chen et al. 2017; Gilad et al. 2018; Musall et al. 2019; Aggarwal et al. 2022).

Despite the prominence of these large-scale dynamics during active wakefulness, the effective role of individual wave events remains an open question. One hypothesis is that waves are largely stochastic events. In this view, their shape and displacement on the cortical surface may passively follow the paths dictated by the network architecture (Ermentrout and Kleinfeld 2001; Mohajerani et al. 2013): they would not be controlled by the subject and would not convey specific information, akin to spike timing in the rate coding interpretation of action potential firing.

Alternatively, individual cortical waves may actively contribute to information processing in the brain. Indeed, cortical state transitions (Crochet and Petersen 2006; Ferezou et al. 2006) and more generally waves (Arieli et al. 1996) directly impact the integration of sensory inputs, leading to either large or very limited evoked activity in response to the same input at the periphery. This contribution of individual waves to information processing is also supported by evidence that they can carry sensory information themselves (Gonzales et al. 2025), and by experiments highlighting that individual cortical waves may be the source or substrate of illusory percepts (Chen et al. 2003; Jancke et al. 2004). However, so far, there has been no direct evidence that individual cortical waves are indeed a relevant functional component of brain activity that could be actively generated by the nervous system towards the resolution of a behavioral task.

The development of motor brain-machine interfacing has emerged as an efficient strategy to interrogate the agency of subjects on internal variables of their brain. In particular, neuronal operant control training has uncovered the ability of a subject to drive the activity of individual neurons (Arduin et al. 2013; Fetz 1969a) as well as small clusters of neurons (Koralek et al. 2012; Clancy et al. 2014; Goueytes et al. 2022) and even pairs of distant neuronal clusters (Clancy and Mrsic-Flogel 2021). Here we ask whether the spatio-temporal dynamics of cortical activity can be orchestrated at the larger scale of cortical waves to meet behavioral needs. In particular, we tested if mice can learn via operant conditioning to generate individual waves with a specific displacement at the surface of the brain. To this end, we monitored in real-time the cortical waves moving across the primary somatosensory cortical area of awake, water-regulated, transgenic mice (Emx1-Cre x Ai-95) expressing the GCaMP6f calcium reporter across excitatory neurons. We tracked online the local maxima of the cortical waves, and we conditioned the delivery of water rewards to the occurrence of wave trajectories that moved through two adjacent 500 µm square patches at the surface of the cortex. Simultaneously, high-speed videography monitored the mouse’s forelimb and hindlimb movements.

We found that a majority of the mice learned to increase the frequency of the Conditioned Waves during training. As they generated cortical waves, the mice deployed a strategy that combined limb movements — which generated excitatory activity in the primary somatosensory cortex — with strong pre-movement cortical state modulations that shaped them into cortical waves that fulfilled our conditioned criteria for reward. This shows that mice can exert a degree of control on the properties of individual activity waves as they address behavioral challenges, and suggests that individual waves are a functionally relevant player of cortical information processing.

## RESULTS

### Mice learned a cortical wave-dependent task

We have developed a novel widefield optical brain–machine interface to test whether mice can learn to generate individual mesoscopic waves with a specific displacement at the surface of the somatosensory cortex in order to get rewards. Sixteen mice expressing GCaMP6f in excitatory neurons of the neocortex were implanted with a head fixation bar and a chronic optical window over the somatosensory and motor cortices (see Materials and Methods). The mice were water-regulated and trained to remain head-fixed on a setup in which they could freely obtain water by licking from a spout. We recorded their spontaneous cortical activity through widefield calcium imaging at 100 Hz (Figure 1A). To detect events of synchronous cortical activity (100 µm-1 mm spatial range), images were filtered with a 2D wavelet transform and the local maxima (x/y coordinates) of these events were tracked in real time at the surface of the cortex. We detected series of consecutive local maxima distant by less than 200 µm. Such a series constituted the trajectory of a dynamic elevation of activity, which we define as a Wave (Figure 1B).

**Figure 1.**
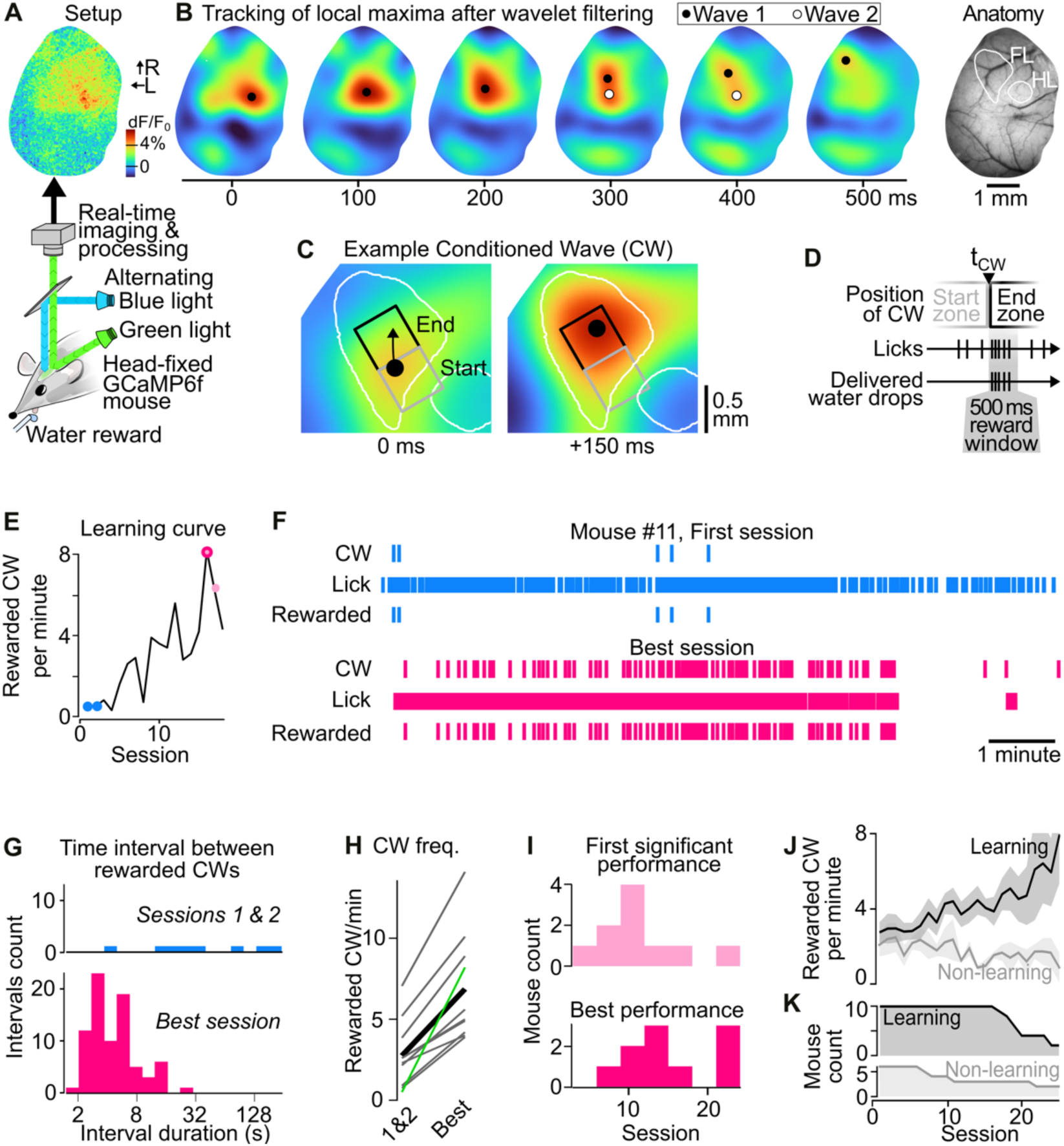
Operant conditioning of spatially defined cortical waves. (**A**) A water-regulated GCaMP6f-expressing mouse with a chronic window above the somatosensory and motor cortices was head-fixed under a macroscope. Alternating imaging with blue/green illumination aimed to correct for intrinsic hemodynamic related signals in functional imaging. Water rewards were conditionally delivered from a spout upon licking. (**B**) Example GCaMP6f fluorescence signals from Mouse #14. Images were acquired at 100 Hz and processed in real-time. After hemodynamics correction, they were filtered with 2D wavelets (100 µm to 1 mm bandpass). The position of several local maxima corresponding to individual Waves could be simultaneously tracked online. Right: Blood vessels at the surface of the cortex shown by green illumination. Hindlimb (HL) and Forelimb (FL) touch representations were functionally mapped by imaging sensory-evoked signals under anesthesia. L, R arrows: lateral and rostral stereotactic orientations. (**C**) In the same mouse, example of a cortical Conditioned Wave (CW): as it travels across the cortical surface, it is detected inside the Start and then the End zone (gray and black squares, respectively, as defined by the experimenter). Black dot: position of the detected local maximum of the wave. White contour: FL and HL touch representations. (**D**) Schematic time course of licks and reward delivery for CWs. t_CW_ is the time at which a CW exits the Start zone and enters the End zone. In the 500 ms window after t_CW_, each lick on the spout triggers the release of a water droplet, collected by the tongue in the same lick movement. (**E**) Frequency of Rewarded CWs across the training sessions of mouse #11. Color code: Blue-filled circle: sessions 1 & 2 used here as reference for untrained state; Light magenta-filled circles: Session where the intervals between consecutive Rewarded CWs were significantly shorter than during sessions 1 & 2. Dark magenta outline: best session (Kruskal-Wallis test followed by multiple comparison tests). (**F**) Example time sequence of CWs, Licks, and Rewarded CWs in mouse #11 during its first session (top, blue) and the session with the highest Rewarded CW frequency (best session, bottom, magenta). (**G**) From the same data as in **F**, histogram of the time intervals between consecutive Rewarded CWs during the first two sessions (top) versus the best session (bottom), shown on a logarithmic scale. (**H**) Magnitude of the difference in rewarded CW frequency between session 1 & 2 and the Best session, across individual mice (gray lines). Thick black line: average. Green line: mouse #11. (**I**) Population distribution of the number of training sessions until the first session with a significant increase in performance (top) and until the best session (bottom). (**J**) Average Rewarded CWs frequency across mice with a significant increase in performance (Learning, black line), versus no significant increase (Non-learning, gray line). Light background: SEM (n values shown in panel **K**). (**K**) Mouse count in the Learning and Non-Learning groups as training progressed.

We selected two nearby 500 × 500 µm square zones (Start and End) within the optical window, and defined Conditioned Waves (CW) as the Waves that were detected first in the Start, and then in the End zone (Figure 1C). The time of first entrance in the End zone t_CW_ initiated a 500 ms opportunity window, during which water droplets were instantly delivered upon licking (Figure 1D). In order to facilitate the initial phase of the learning, we selected the location of the Start and End zones so that 1 to 5 Conditioned Waves occurred per minute during spontaneous brain activity. Because of the diversity of local wave statistics observed across mice, the location of the Start and End zones varied within the somatosensory cortex, in the vicinity of the limb areas, that we identified by recording GCaMP6f signals evoked by passive limb stimulations under anesthesia (Figure 1B,C, Material and Methods).

We trained the mice daily, during sessions lasting 20 to 30 minutes. We quantified performance — defined as the frequency of Rewarded CWs — and carried out all subsequent analysis on the first 10 minutes, when the highest motivation was apparent (Matteucci et al. 2022). To assess the evolution of mice performance with training (example in Figure 1E), we computed for each session the distribution of the time intervals between Rewarded CWs, and compared it to sessions 1 and 2 combined (Figure 1F,G, Methods). Of the 16 trained mice, 10 increased significantly their performance compared to the first 2 sessions ("Learning" group, example outcome from Kruskal-Wallis test followed by multiple comparison tests shown in Figure 1E). In the Learning group, the averaged frequency of Rewarded CWs was 2.5 times higher in the best session compared to sessions 1 & 2 (Figure 1H). Their first significant performance was achieved after 11.1 ± 5.4 training sessions (mean ± SD, Figure 1I, top). Several of these mice kept progressing with additional training, so that on average the best performance was reached after 15.0 ± 5.6 training sessions (Figure 1I bottom, Figure 1J). It should be noted that although not all the mice underwent the same number of training sessions, all Learning mice were trained at least during 18 sessions (Figure 1K).

### A selective increase in frequency of Conditioned Waves with training

The increase in performance observed in the Learning group may be due to a change in cortical dynamics and/or of the licking behavior. The quantification of detected waves revealed a significant increase in the frequency of CWs (Figure 2A) in the context of an overall increase in Wave frequency within the imaged window (Figure 2B). We asked if these learning-driven changes were caused by a homogeneous increase in Wave frequency across the cortical surface, or if instead the mice managed to generate a spatially focused increase of CWs frequency. To probe this, we explored the spatial structure of the Wave trajectories, and more specifically the spatial density of their individual time points (examples for Sessions 1 & 2 in Figures 2C,D, Best session in Figure 2E). With learning, this wave density increased mainly around the Start/End zones (Figure 2F, population analysis in Figure 2G).

**Figure 2.**
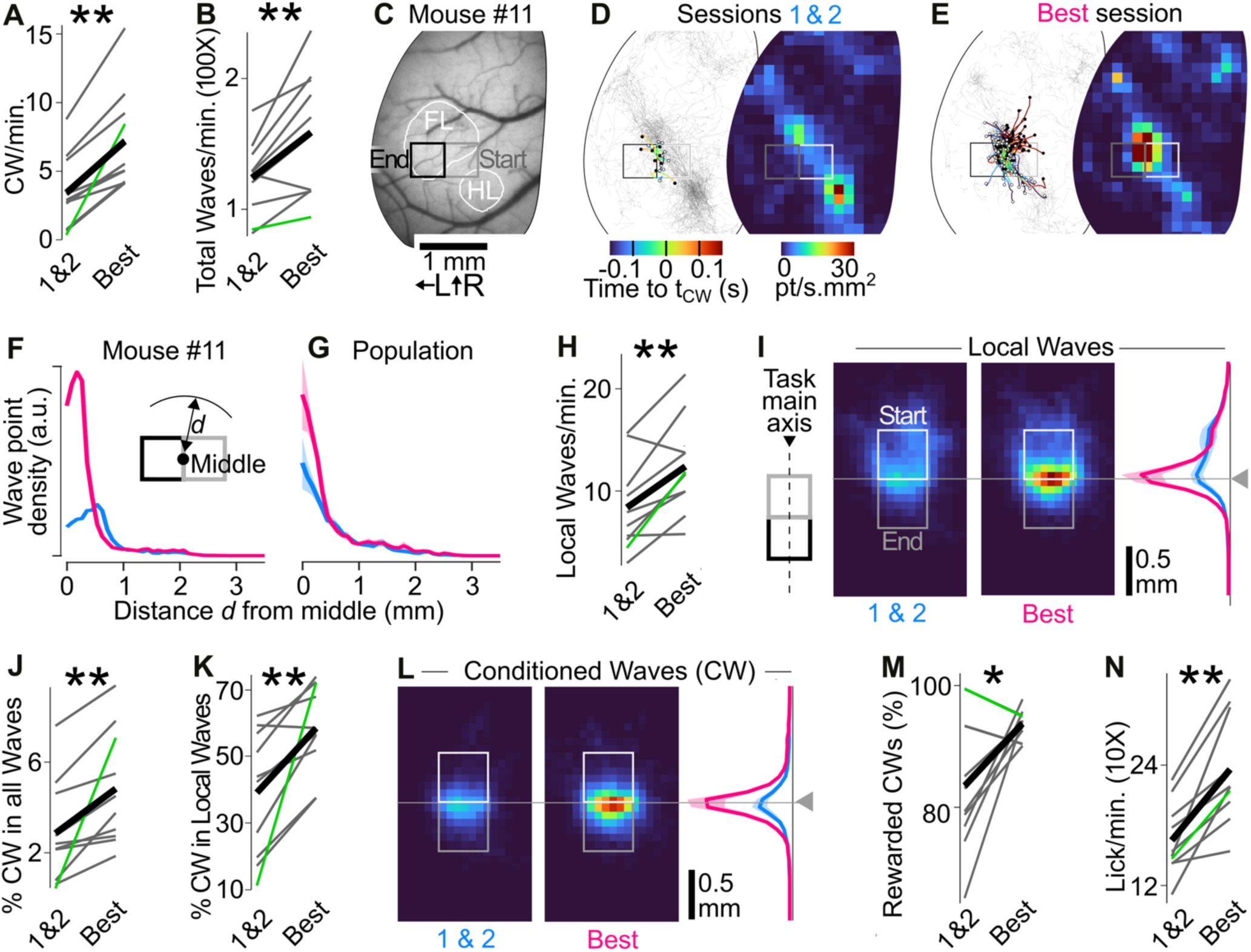
Learning is dominated by a selective increase in frequency of Conditioned Waves. (**A**) Frequency of Conditioned Waves (CWs) during the first two sessions and during the session with the best performance. **: Wilcoxon p = 0.002. Gray: individual mice. Green: mouse #11. Black: population average. (**B**) Same as **A** for the frequency of all Waves detected on the imaged cortical surface. **: Wilcoxon p = 0.0098. (**C**) Case study: Location of the Start and End zones for example mouse #11. White delineations: primary FL and HL areas of the cortex identified by passive stimulations (see methods). L, R arrows: lateral and rostral stereotactic orientations. (**D**) Left: Trajectories of all Waves detected during sessions 1 and 2 in mouse #11 (gray), overlaid with trajectories of CWs (colored). The changing color along CW trajectories indicates time relative to t_CW_. White dot: first point of CW. Black dot: last. Right: point density of all Wave trajectories. (**E**) Same as **D** for the best session. (**F**) Radial distribution n of Wave point density, computed for all Waves of mouse #11. First and second (blue) versus best session (magenta), centered on the middle point of the task (see inset schematic). (**G**) Same as F at the population level in the Learning group. Light background: SEM. (**H**) Evolution of the frequency of Local Waves: all Waves detected in the Start zone, including CWs. **: Wilcoxon p = 0.0059. (**I**) Population average of the density of the Local Wave points at the surface of the cortex, around the Start and End zones. Left: Sessions 1 & 2. Middle: Best session. Right: Marginal distribution of Local wave point density along the task main axis, which sets the position and alignment of the Start and End zones. Colormap: same as in **D**. (**J**) Evolution of the proportion of CWs among all recorded Waves. **: Wilcoxon p = 0.002. (**K**) Same as **A** for the percentage of CWs among Local Waves. **: Wilcoxon p = 0.0039. (**L**) Same as **I** for CW points. (**M**) Evolution of the proportion of Conditioned Waves that were actually rewarded. *: Wilcoxon p = 0.027. (**N**) Evolution of the frequency of licking. **: Wilcoxon p = 0.002.

The occurrence of Local Waves — defined as all Waves that were detected in the Start zone, even if not reaching the End zone — increased significantly (Figure 2H). Following learning, we found that they spent most of their duration at the interface between the Start and End zones (Figure 2I). When we then focused on CWs, we found that their proportion increased both with respect to all Waves (Figure 2J) and with respect to Local Waves (Figure 2K). Consistent with the general trend observed in Local Waves, learning led CWs to become spatially focused on the interface between the Start and End zones (Figure 2L). Overall, training led to a specific increase in the frequency of cortical waves that were spatially positioned and oriented to match the operant conditioning rules we set.

Among CWs, a large and significantly increasing proportion was rewarded (Figure 2M). This came together with a strong increase in the frequency of licking (Figure 2N), suggesting that the mice learned to continuously lick in order to take advantage of any reward opportunity, rather than coordinate licking with CWs (but see Figure 3). This observation led us to focus the rest of our analysis on the timing of Conditioned Waves t_CW_, which are at the core of task learning and constitute a good proxy of the rewards obtained by the mice.

**Figure 3.**
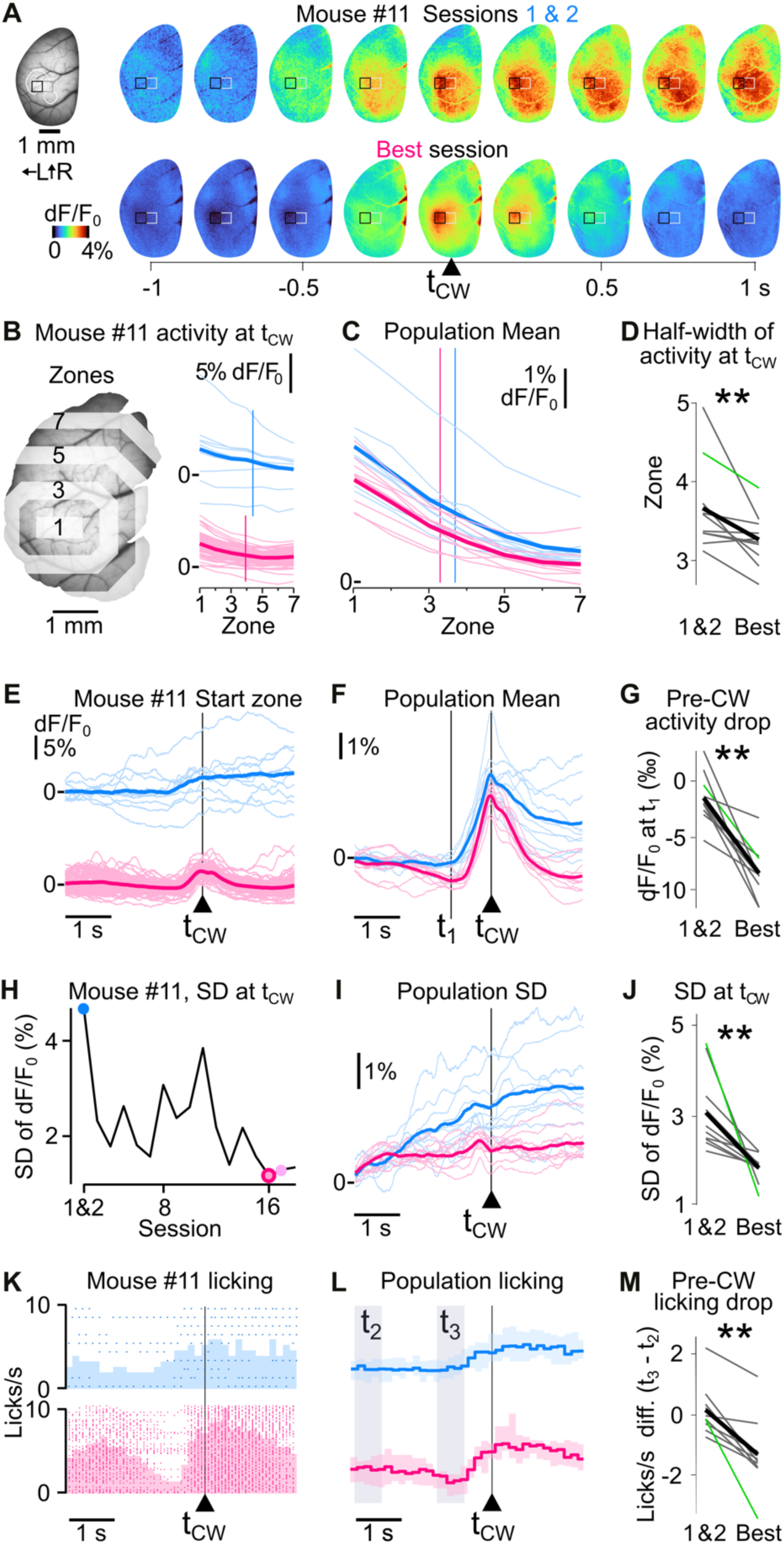
Spatiotemporal shaping of cortical dynamics by the operant conditioning of Wave trajectories. (**A**) Average GCaMP6f signal across the cortical window in mouse #11, aligned on t_CW_, during sessions 1 & 2 (top, n = 10 waves) and the best performance session (bottom, n = 86). L, R arrows: lateral and rostral stereotactic orientations. (**B**) Calcium fluorescence signal averaged at t_CW_ in the spatial zones 1-7 located at increasing distances from Start and End zone combined (zone 1); same mouse as in **A**. Thin lines: individual CWs. Thick line: average across all CWs of the session. Top, blue: sessions 1 & 2. Bottom, magenta: best session. Vertical lines: spatial extent of calcium signal quantified as distance at half maximum amplitude (half-width). (**C**) Population average of the calcium fluorescence signal at t_CW_ in the spatial zones 1-7. Thin lines: individual mice. Thick line: population average. Blue: sessions 1 & 2. Magenta: best session. Vertical lines: population average of half-width of calcium signal spatial spread during sessions 1 & 2 (blue) and the best performance session (magenta). (**D**) Population analysis of spread (half-width) of calcium activity at t_CW_. **: Wilcoxon p = 0.006. Gray: individual mice. Green: mouse #11. Black: population average. (**E**) Calcium fluorescence signal averaged in the Start zone aligned on t_CW_ from the same mouse as in **A**. Thin lines: individual CWs. Thick line: average across all CWs of the session. Top, blue: sessions 1 & 2. Bottom, magenta: best session. (**F**) Population average of the calcium fluorescence signal around t_CW_. Thin lines: individual mice. Thick line: population average. Blue: sessions 1 & 2. Magenta: best session. (**G**) Population analysis of the pre-CW suppression (quantified at the lowest point indicated as t_1_ in **F**). **: Wilcoxon p = 0.002. Gray: individual mice. Green: mouse #11. Black: population average. (**H**) Evolution of the standard deviation of the ‘Start’ zone calcium fluorescence signal across individual CWs, quantified at t_CW_, throughout sessions for mouse #11. Blue-filled circle: sessions 1 & 2. Light magenta-filled circle: sessions with significantly higher Rewarded CW frequency as in sessions 1 & 2. Dark magenta outline: best session. (**I**) Time course of the standard deviation across individual CWs, aligned on t_CW_. Thin lines: individual mice. Thick line: population average. (**J**) Population analysis of the evolution of the standard deviation of the ‘Start’ zone calcium fluorescence signal across individual CWs, quantified at t_CW_, **: Wilcoxon p = 0.002. (**K**) Distribution of licks around t_CW_ in the first two (top) versus best session (bottom) in mouse #11. The raster of licking times is shown on top of the time histogram. (**L**) Averaged instantaneous licking frequency aligned on t_CW_ during sessions 1 & 2 (blue) versus the best session (magenta). Light background: SEM. Y scale: same as in K. (**M**) Change of lick rate quantified between t_2_ and t_3_ as defined in **L**. **: Wilcoxon p = 0.002.

### Spatiotemporal shaping of cortical activity and licking around CWs

So far, we focused our analysis on the evolution of the trajectories of Waves that were specifically reinforced by operant conditioning. We next asked if the underlying mesoscale cortical dynamics was also shaped during this learning process. To this end, we analyzed the averaged, hemodynamics-corrected, non-filtered widefield calcium signals acquired around each t_CW_. During the first two sessions (combined), the calcium activity around t_CW_ exhibited a broad spatial extent (example in Figure 3A top, population mean in Figure 3B,C blue). In contrast, after training, the average calcium activity around CW time was more focused in space around the Start/End zones (example in Figure 3A bottom, population mean in Figure 3B,C magenta), and there was a significant reduction in half-width of the spatial extent of activity at t_CW_ (Figure 3D). In parallel to this focusing of the CW-associated cortical activity, a strong suppression became visible with training, both before and after the CWs (example in Figure 3A, E, Supplementary Movie 1). The drop of activity just before the CW was significant at the population level (Figure 3F,G). In addition, we found that the inter-trial variability in calcium signal dynamics decreased with training (example in Figure 3H, population analysis in Figure 3I,J). This reduced variability suggests that mesoscale cortical dynamics — and in particular the pre-CW suppression — were operantly-conditioned and might therefore be required for the initiation of the cortical Waves that were rewarded in our task.

Remarkably, we found that licking was also significantly down-regulated at the same time as mesoscale cortical activity (Figure 3K-M) after learning, on top of an overall increase in licking that we described earlier (Figure 2N, also visible in Figure 3K).

### Movement and cortical dynamics combine to generate Conditioned Waves

Next, we asked what was the origin of the coordinated cortical activation that is generated by the mice as they solve the task. In the limb areas of the primary somatosensory areas where we positioned the Start and End areas, a major source of activity should be the limbs peripheral somatosensory inputs (proprioceptive and touch inputs) as well as additional movement-related activity, including motor inputs from higher areas. Therefore, we hypothesized that during the task, the mice may perform limb movements to trigger cortical activity towards solving the task. To probe this hypothesis, we tracked the right forelimb and hindlimb position through all sessions (for 8 out of 10 ‘Learning mice’, see Methods). We averaged the speed of the two limbs into an overall “limb speed” variable (example in Figure 4A) that we used to detect individual limb movements based on a speed threshold (speed > 15 mm/s, see Methods). We often found — even during the best performance session of trained animals — repeated limb movements that persisted until one was immediately followed by a CW (Figure 4A, movement examples in Figure 4B, associated cortical waves in Figure 4C, time courses in Figure 4D). Overall, although some CWs were not associated with any movement (Wave 6 in Figure 4A-D), limb movements featured consistently before CWs (Figure 4E). At the population level, 92.4% of the CWs during the first two sessions were immediately preceded by limb movements, a proportion that even increased slightly with learning (Figure 4F).

**Figure 4.**
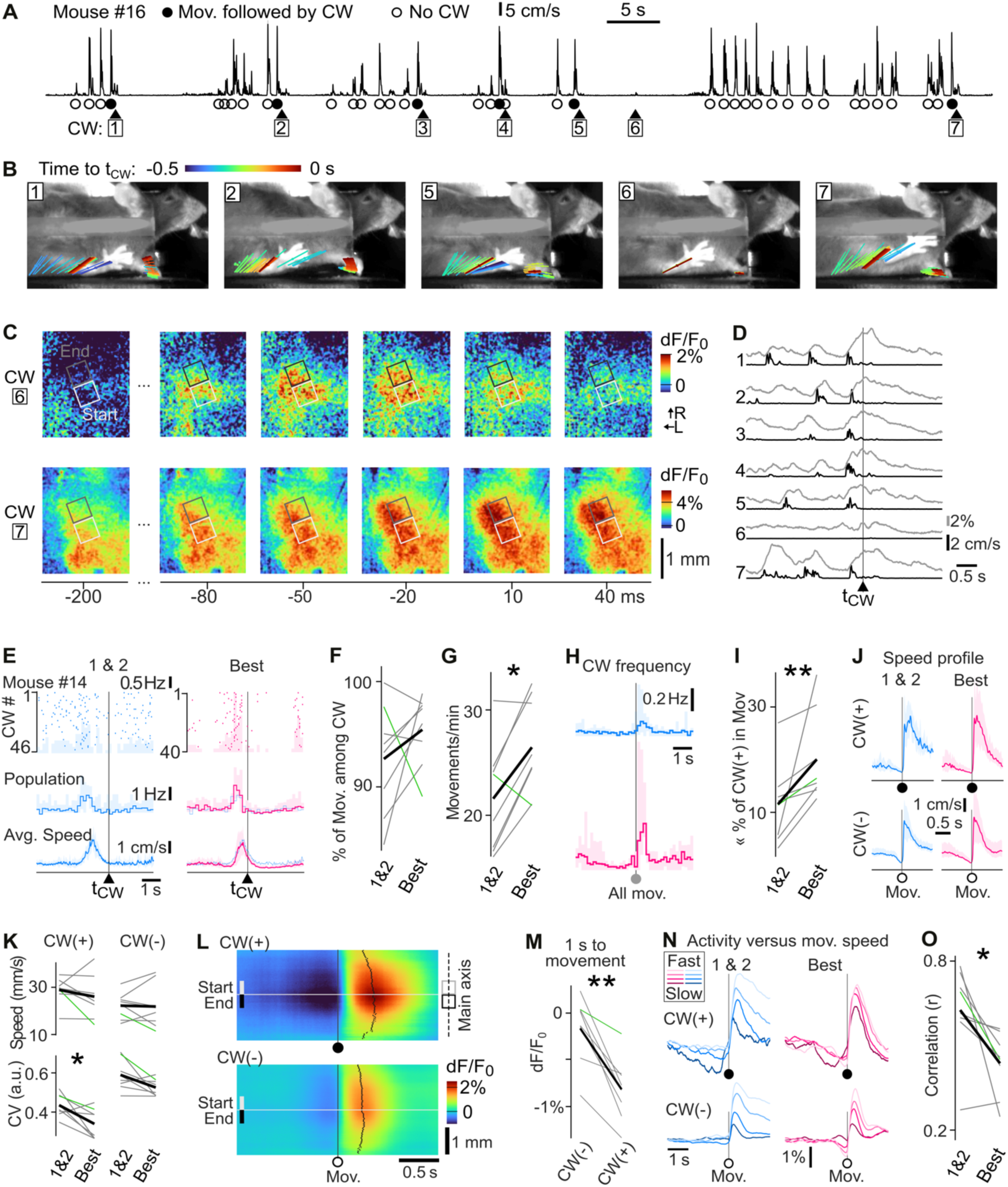
Conditioned waves stem from the interaction between forelimb movement activity and local cortical state modulation. (**A**) Limb speed (average of forelimb and hindlimb) of mouse #16, during 90 s of the best performance session. Triangles: CWs 1 to 7. Filled circles: movements associated with a CW, that is, when t_CW_ is between movement onset and 100 ms after movement offset. Open circles: movement without a CW. (**B**) Side view of mouse #16 behavior, overlaid with the movements of the right hindlimb and forelimb palms in a 0.5 s window prior to t_CW_ in 5 of the CWs shown in **A**. (**C**) Snapshots of the calcium signal imaged around t_CW_ for CWs 6 and 7 (see **A** and **B**). White: Start zone. Black/Gray: End zone. (**D**) Superimposed limb movements (black) and average calcium dynamics quantified over the Start zone (gray) around the 7 CWs shown in **A**. (**E**) Distribution of the limb movements around t_CW_, in the first two sessions (left, blue) and the best session (right, magenta). Top: scatter plot of detected limb movement onsets and corresponding time histogram for all CWs of mouse #14. Middle: population median of the limb movement onsets time histogram (n = 8). Light background: 5% and 95% percentiles. Bottom: Average limb speed. Light background: standard deviation. Population data from sessions 1 & 2 is repeated in the background of plots for the best session data. (**F**) Evolution of the proportion of CWs that were preceded by a detected movement onset between the first two sessions, and the best session. Green line: mouse #16, thick line: population average. (**G**) Evolution of the frequency of detected limb movements *: Wilcoxon p = 0.039. (**H**) Population median of the time distribution of the t_CW_ around all movement onsets. Light background: 5% and 95% percentiles (n = 8). (**I**) Proportion of the movements that are immediately followed by a CW (latency ≤100 ms after movement offset). **: Wilcoxon p =0.008. (**J**) Population average speed profile of the detected movements in the first two sessions (left, blue) and best session (right, magenta). n = 8. Top: movements associated with a CW (CW(+), filled circle: movement-onset). Bottom: other movements (CW(-): open circle). Light background: standard deviation. (**K**) Average and coefficient of variation (CV) of the peak speed of the CW(+) and CW(-) movements across learning (sessions 1 & 2 versus best session). The coefficient of variation decreased significantly for Conditioned Waves. *: Wilcoxon p = 0.023 (n = 8). (**L**) Population average cortical dynamics around movement onsets across the task main axis, for CW(+) (top) and CW(-) movements (bottom), in the best session (n = 8). Black curve: time of the peak of the activity along the task main axis. (**M**) Population average of the pre-movement calcium signal corresponding to the CW(+) and CW(-) movements during the best session. The signal was averaged in space in Start and End zones and in time in one second preceding the movement onset. **: Wilcoxon p = 0.008 (**N**) Evolution of the population average calcium dynamics in the Start zone, aligned on movement onsets, split by movement speed. Left: first two sessions. Right: best session. (**O**) Population evolution with training of the Pearson correlation coefficient between the peak activity in the Start zone (t_CW_ - 0.1 to t_CW_ + 0.5 s) and the average limb movement speed between t_CW_ - 1.5 s and t_CW_ + 0.5 s. *: Wilcoxon p = 0.016.

However, even if the frequency of the limb movements irrespective of Conditioned Waves increased significantly (Figure 4G) and CWs were increasingly aligned on movements (Figure 4H,I, see also Figure 4E), the proportion of all movements that were followed by a CW remained low (20%, Figure 4I). These frequent failures of limb movements to generate CW may be explained by inadequate or inconsistent movements. The comparison of the profile of the limb movements that led to CW (labelled CW(+) movements) versus limb movements that did not lead to a CW (CW(-)) did not reveal any clear distinctive feature. The average movement speed time course remained similar (Figure 4J), while we observed a specific and significant reduction of the variability of the CW(+) movements (Figure 4K). Overall, this suggests that the mice did stabilize and reinforce a subset of CW-leading movements, although still ∼80% of the movements were CW(-) at the end of the training period.

Stereotyped limb movements are therefore unlikely to be the only necessary requirements to enable the emergence of a CW. We hypothesize that in addition to shaping their limb movements to generate peripheral inputs that would yield a Conditioned Wave in the cortex, the mice combined the large cortical activation triggered by peripheral movements with an active sculpting of these inputs by cortical state dynamics.

To probe this, we compared the cortical dynamics associated with CW(+) versus CW(-) movements. Despite comparable movement in these two conditions, we found very different cortical dynamics. In particular, in trained mice, before CW(+) movements, we found a strong and significant suppression of activity that was absent before CW(-) movements (Figure 4L,M). In addition, the magnitude of the activation following the movement was higher for CW(+) movements (Figure 4 L,N).

To understand if this difference in cortical state before movement may have had an impact on the cortical dynamics triggered by the limb movements, we sorted them by limb speed. In the first training sessions, there was a direct link between limb speed and calcium wave amplitude, for both CW(+) and CW(-) movements (Figure 4N, left). However, in the best session, after training, this relation appeared largely abolished in CW(+) movements, but not in CW(-) movements (Figure 4N, right). Consistently, computing the linear regression between the speed of individual movements and amplitude of the corresponding calcium activation confirmed a correlation that was significantly reduced with learning (Figure 4O, Wilcoxon p = 0.016).

Overall, these findings show that with learning, the production of Conditioned Waves remained largely tied to peripheral movements. However, we found that these cortical activations were shaped by pre-movement cortical dynamics into CW, and became increasingly independent from key limb movement characteristics such as speed.

### Spatial separation of Start and End zones stretches cortical dynamics

After training on the initial protocol during 21 sessions on average, we aimed to probe the ability of the mice to generate longer cortical waves. To this end, we transitioned the mice to a Gap condition, where the Start and End zones were separated by 130 µm. To solve this new task, the mice had to produce a Long Wave that crossed this space gap in order to obtain a reward (Figure 5A). We maintained the rest of the protocol unchanged. Nine mice were trained with this Gap protocol for an additional 13 sessions on average.

**Figure 5.**
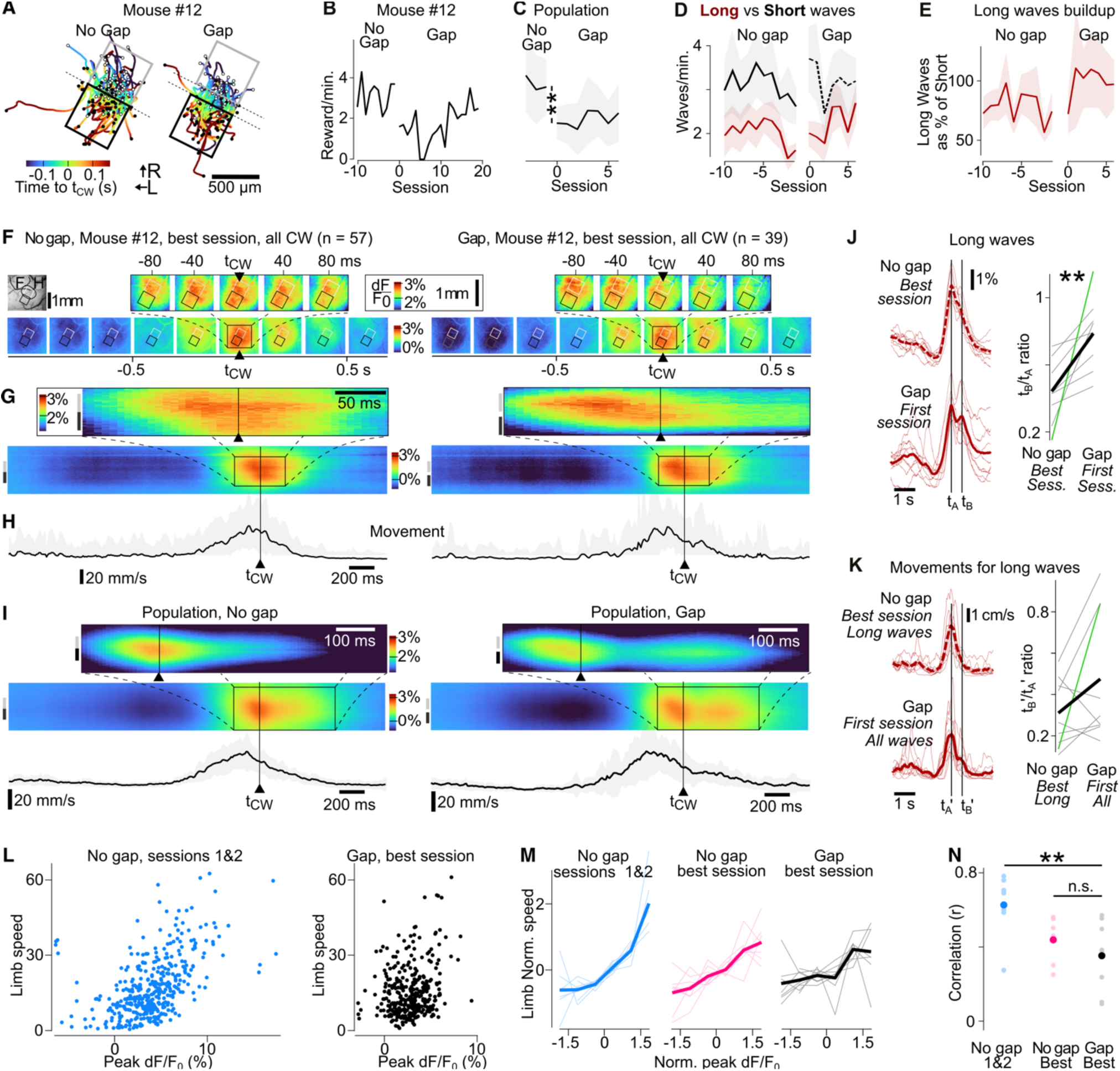
The operant conditioning of cortical waves travelling longer triggers additional reshaping of cortical dynamics. (**A**) Example trajectories of the detected CWs in the No Gap (left) versus the Gap condition (right) in mouse #12 (best session). Color code: time around t_CW_. (**B**) Example performance (Rewarded CW frequency) of mouse #12 around the transition from the No-Gap to the Gap condition. (**C**) Same as **B** at the population level. Light background: SEM. *: Wilcoxon p = 0.0080 (n = 9 mice). (**D**) Evolution of the frequency of Waves that would fulfill the criteria of the Gap condition (Long Waves, brown), and of Waves that would fulfill the No Gap condition, but not the Gap condition (Short Waves, black). Continuous lines: fulfill the ongoing task condition. Light background: SEM (n = 9 mice). (**E**) Ratio between the count of Short and Long Waves. Light background: SEM (n = 9 mice). (**F**) Example in mouse #12 of the average of t_CW_-aligned cortical dynamics during the initial, No Gap condition (left, best session in the condition) versus the Gap condition (right, best session in the condition). (**G**) Time/Space view of the same data as in **F**. Data was projected on the task main axis. (**H**) Average limb speed around t_CW_, No-Gap (left) versus Gap (right) condition, in the same mouse example. Light background: SEM. (**I**) Population analysis matching **G**-**H** (n = 9 mice). (**J**) Left: population average time profile of the calcium signal in the Start zone during Long Waves that match the Gap condition, in the best No Gap session (Top), versus the first Gap session (bottom). The waves are time-aligned on their maximum. Right: Ratio extracted from the Long Waves, between the time of the wave peak (t_A_) and the time of the rebound observed in the Gap condition (t_B_, 0.5 s after t_A_). Green line: mouse #12. **: Wilcoxon p = 0.0040. (**K**) Same as **J** for limb movements, also aligned on the peak of the cortical activity in the Start zone. t_A_’ and t_B_’ are shifted compared to t_A_ and t_B_ so that t_A_’ is at the peak of the limb movements average. We found no significant difference. (**L**) Peak value of the cortical activity versus average limb speed around t_CW_. The data points of all mice are merged. Left (blue): No gap condition, sessions 1&2. Right (black): Gap condition, best session. (**M**) Average normalized (reduced and centered) Limb speed observed before t_CW_ as a function of the average cortical activation in the Start zone at t_CW_. Thin lines: individual mice. Thick lines: population averages. (**N**) Pearson correlation between the CW-related cortical activity and movements, as reported in **M**. **: Mann-Whitney p = 0.002.

At the time of the transition from the No gap to the Gap protocol, the frequency of Rewarded CWs dropped significantly (Figure 5B,C). This was expected since through the No gap training, the frequency of Long Waves matching the requirement of the Gap protocol was lower than that of Short Waves whose trajectory only fulfilled the No gap task requirements (Figure 5D). This lower frequency of Conditioned Waves persisted throughout the training in the Gap protocol. Still, we found that the ratio of Long Waves over Short Waves, which was stable in the No gap condition, increased, although not significantly, with training in the Gap condition (Figure 5E).

Overall, this Gap protocol was more challenging than the No gap condition for the mice and on average they did not manage to recover a large Rewarded CW frequency (Figure 5C), although some mice were successful at increasing performance in the Gap condition (example in Figure 5B).

The mice performed this new task by building on their ability to solve the initial No gap protocol, and further reshaped their cortical dynamics. In particular, they maintained the cortical hyperpolarization before wave onset (Figure 5F-I, No gap: left, versus Gap: right column), and generated more extended CWs when compared to the No gap condition (Figure 5J). These extended CWs appeared to split into a first main wave followed by a significant rebound of activity (Figure 5I,J). Except in 3 mice, this second phase of the Gap CWs was not associated with a matching evolution of limb movements (Figure 5K).

To further probe this dissociation of the CW dynamics from the associated movements, we repeated our study of the link between limb movement speed and the amplitude of the cortical dynamics, which we found was already diminished by learning in the No gap condition (Figure 4L-N). A scatter plot of all mean limb speed around t_CW_ versus the peak calcium dynamics across the whole population showed that in contrast to the clear positive relationship observed in non-trained mice (Figure 5L-N Pearson r = 0.63), the linear relationship was lost after the training, first in the No gap, and then in the Gap protocol (Figure 5L-N, r = 0.35 in the best Gap session). Overall, this clear loss of the link between movement and calcium activity in the primary somatosensory cortex was even more visible in the No gap than in the Gap condition (Figure 5N). In summary, in attempting to generate these more challenging mesoscale dynamics to address the Gap condition, the mice used the same strategy as for the No gap condition, but they amplified the cortical shaping of the movement-related activation as they generated an increased proportion of long waves. The resulting cortical waves displayed unexpected features, including a rebound of activity after t_CW_.

## DISCUSSION

Using an optical brain-machine interface based on the real-time processing of wide field calcium signals, we have shown that mice can generate specific cortical waves of neuronal activity at the mesoscopic scale in order to solve an operant conditioning task. From the beginning of the training, Waves matching the imposed criteria of the task appeared immediately after a limb movement, indicating that part of the underlying activity could rely on limb movement-related cortical activity. In naïve animals, we observed a correlation between Conditioned Wave amplitude and limb movement speed, but this link disappeared as training progresses. Remarkably, the vast majority of limb movements did not generate Conditioned Waves, even in trained mice. Only movements occurring after a local suppression of cortical activity led to the emergence of Conditioned Wave. This suppression of cortical activity preceding Conditioned Waves, which increases significantly with training, appeared to have a key role in cortical wave shaping. Finally, when the task was made more difficult by asking the mouse to generate Waves with extended trajectories, we observed a reshaping of the Conditioned Waves. It was associated with the emergence of a rebound of activity, while the link between movement and activity continued to decrease. Altogether these finding suggest that mice could shape their cortical activity at the mesoscopic scale in a goal-directed manner, by taking advantage and reshaping the physiological cortical activation associated with limb movements.

### A learning-based mesoscale brain-machine interface

Invasive brain-machine interfacing provides a unique opportunity to probe the information processing architecture of the brain, by extending the operant conditioning paradigm to internal variables of the brain. However, so far, strategies based on learning have only been successful for small groups of neurons (Arduin et al. 2013; 2014; Clancy et al. 2014; Fetz 1969b; Koralek et al. 2012; Neely et al. 2018; Prsa et al. 2017), small (∼0.1 mm^2^) patches of cortical tissue trained through the real-time processing of local 1-photon calcium imaging (Clancy and Mrsic-Flogel 2021), and LFP recordings (Engelhard et al. 2013; Shi et al. 2024; Chauvière and Singer 2019). Later experiments showed that when a group of neurons was conditioned to solve a task, a single neuron often took the lead in controlling the brain-machine interface (Abbasi et al. 2023; Jeon et al. 2022). Here, in contrast, we conditioned the delivery of a reward to the coordinated activity of a large population of cortical neurons shaped into a traveling peak of mesoscopic activity. Therefore, our experiments open up the possibility to operantly-condition the spatio-temporal features of neuronal activity at a scale that is relevant to electrocorticogram measurements (ECoG). Interestingly, the ECoG signals have already been exploited for motor decoding strategies in humans, both decoding of speech (Anumanchipalli et al. 2019), and body control restoration (Lorach et al. 2023). Our experiments also revealed the buildup of large-scale changes in activity across several millimeters of the imaged cortical area, both before and after the CW (Figure 3,4 and 5). We hypothesize that these indirectly conditioned changes of the cortical activity are the sign of fundamental constraints on the plasticity of cortical mesoscale dynamics.

### A simple readout of complex mesoscale cortical dynamics

In the context of our brain-machine interface experiments, we trained mice to control parameters of individual cortical mesoscale waves. These waves show complex and variable shapes that can be hard to capture through a limited set of descriptors. In fact, the identification of such descriptors has been a major effort in the field of optical imaging. One strategy has been to identify principal components of the spatial dimensions of the imaged cortical dynamics, and to project back the spontaneous spatiotemporal dynamics into the identified subspace (Mohajerani et al. 2013; Musall et al. 2019). Another strategy has been to compute the flow fields of wave speeds at the surface of the cortex, and focus on the singularities that structure these fields, and in particular sources and sinks (Liang et al. 2023; Mohajerani et al. 2013).

Given the challenge of operant conditioning, which requires a simple variable that could be computed in real-time and manipulated by the mouse, we selected a first order descriptor: the wave position. Several definitions of wave position are possible, including the position of the local maximum within wave territory, as well as the position of the wavefront. However, the wavelet filtering of the raw signal that is required to extract the dynamics of individual waves in real-time (Figure 1) precluded an accurate measurement of the wavefront. Therefore, we chose to track the local maximum of the wave, as this measure is less affected by filtering artefacts. This means that the constraints that we applied to the cortex were local to the ∼500 µm area at the Start/End zones border, while the mice deployed mesoscale dynamics at a millimetric scale in order to solve the task (Figures 3-5).

### Mesoscale cortical activations rely on peripheral movements

As the mice learned the task, Conditioned Waves were mainly generated in conjunction with limb movements (Figure 4F). We hypothesize that touch and proprioception generated from these movements, as well as motor commands in the nearby limb motor cortex, all contributed to the activation of the limb-related area of the somatosensory cortex, which is the location where we have positioned our operant-conditioning windows. It is therefore likely that the mechanism used by the mice to generate the bulk activation of cortical tissue required by the task relied on the cortical mesoscale activation that can be triggered by sensory inputs (Jancke et al. 2004; Ferezou et al. 2007; Afrashteh et al. 2021). This does not match the lack of movement reported in several brain-machine interfacing tasks that relied on the conditioning of few cortical neurons (Clancy et al. 2014; Clancy and Mrsic-Flogel 2021; Prsa et al. 2017). We hypothesize that this is because the bulk cortical activation that is required in our task can be hard to generate for the cortical circuitry in the absence of peripheral inputs, in contrast to the activation of a small subset of neurons, that may be achieved locally by the cortico-striatal circuitry (Koralek et al. 2012) without the need for peripheral inputs.

### Interplay of peripheral inputs with local cortical state

In our experiment, only a small minority (20%) of the movement-triggered bulk cortical activation resulted in a reward (Figure 4I), and the movements that led to either successful or unsuccessful generation of Conditioned Waves were similar (Figure 4J,K). In contrast, the cortical dynamics were vastly different. In particular, we observed a pre-wave suppression of cortical activity that was specific to the Conditioned Waves (Figure 4L), and became stronger with learning (Figure 3E-G). Furthermore, while limb movement amplitude was correlated with the magnitude of associated cortical activation in naïve mice, we observed a specific suppression of this relationship for the movements preceding a Conditioned Wave throughout learning (Figure 4N,O, Figure 5L-N). These findings suggest that the mice actively modulated self-generated somatosensory inputs using the internal cortical dynamics that were reinforced with operant conditioning. This internal processing was likely hard to pull out for the mice, as the percentage of movement associated with Conditioned Waves remained low (despite learning, Figure 4I), and depended on the features of the cortical wave that had to be generated, as shown by the drop of performance in the Gap condition (Figure 5B). The rebound in activity observed after Conditioned Waves in the Gap task configuration (Figure 5J,K) suggests that orchestrating the neuronal dynamics required by the new rule demands an even more complex form of adaptation that calls for further investigation.

### Relationship of Conditioned Waves with spontaneous waves

Several of our findings suggest that to solve the behavioral task, the mice were able to control the timing and adjust the parameters of existing wave patterns, but that they were not able to generate new wave patterns from scratch. In particular, we have observed that learning was facilitated when the selected Start and End zones were located within a cortical region that spontaneously features a high wave density, in our case in the primary somatosensory areas (Figure 2C-D). In addition, the Conditioned Waves retained the local orientation observed in the first sessions through the training sessions, even when this orientation was not optimal to achieve the task (Figure 2D,E). Finally, we have found that it was challenging to extend the distance traveled by the Conditioned Waves beyond their naturally occurring size (Figure 5B-E)

Therefore, Conditioned Waves in our experiments are likely to rely on a similar anatomical substrate as spontaneous cortical waves. Still, these waves were different from spontaneous waves: they were preceded by a large suppression of activity after training; they lost their relationship to limb movement amplitude, and they were followed by activity rebounds in the Gap condition.

### Suppression of the activity around Conditioned Waves

One of the most striking difference with spontaneous waves is the build-up with learning of a suppressed cortical state that precedes and follows Conditioned Waves (Figure 3E-G). This suppression of ongoing activity is reminiscent of the quiet cortical state that has been well described in the somatosensory cortex (Petersen et al. 2003; Crochet and Petersen 2006; Ferezou et al. 2007). A transition to a quiet state in preparation for the generation of a cortical wave triggered by peripheral inputs would optimize its magnitude, as sensory inputs trigger stronger cortical activations in this context than during active cortical states (Fanselow and Nicolelis 1999; Petersen et al. 2003; Crochet and Petersen 2006; Ferezou et al. 2007). However, in several ways, the dynamics we observed around Conditioned Waves do not match the established features of the typically reported “quiet” state. In particular, we observed that after learning, the Conditioned Waves were shortened and followed by a second phase of hyperpolarization (Figure 3F) while sensory inputs in the context of a quiet state should have transitioned the cortex to a long-lasting active state, rather than rapidly reverse to a quiet state. In addition, we found that after learning, the magnitude of the Rewarded Waves was not anymore modulated by the amplitude of the mouse movement, in contrast to the established observations of cortical integration of inputs with cortical state (Petersen et al. 2003; Crochet 2011). Finally, we found that this transitory suppression of activity was time-locked to CW-related movement onsets (Figure 4L,M) and was accompanied with a structured modulation of the licking (Figure 3K-M).

This suggests that the pre-Conditioned Wave suppression is part of a voluntary action planning for the generation of a Conditioned Wave, rather than a transition to quiet state, which is generally associated with non-attentive, non-task engaged behavior. In particular, it could be related to an attentional mechanism, similar to the attention-driven spatial focus in the primary visual area, where non-task related activity is dampened, in contrast to task-relevant inputs (Brefczynski and DeYoe 1999; Dugué et al. 2020; Herrmann et al. 2010; Murray 2008). In visual tasks, such attentional effects have been linked to higher order cortical areas such as the frontal eye field, as well as to the pulvinar and thalamic nuclei.

Overall, we conclude from these observations that the mice generated a specific cortical processing state in order to generate Conditioned Waves from an existing source of bulk cortical activation. This remained challenging even after repeated training, as it was only successful in 20% of the limb movements (Figure 4I). Previous work on BMIs based on the readout of individual neurons have shown that the cortico-striatal loop is required to enable the learning of a new pattern of single neuron activity in order to solve a BMI task (Koralek et al. 2012; Neely et al. 2018). Further experimental work will be required to determine the actual subcortical circuitry and mechanisms that support the generation of Conditioned Waves in our experiments, including the suppression of cortical activity prior to Conditioned Waves.

## MATERIALS AND METHODS

### Animals

Experiments were performed on adult Emx1-Cre (B6.129S2-Emx1^tm1(cre)Krj^/J; Jax # 5628) x Ai95 (B6;129S-Gt(ROSA)26Sor^tm95.1(CAG-GCaMP6f)Hze^/J; Jax # 24105) transgenic mice expressing GCaMP6f in cortical excitatory neurons (Madisen et al., 2010). Male and female animals were used for the experiments (n = 16: 11 males, 5 females). Housing was enriched with a wheel, a tunnel, nesting material, and toys in a 12-hour light cycle, with food *ad libitum*. Protocols were in accordance with the French and European (2010/63/UE) legislations relative to the protection of animals used for experimental and other scientific purposes. All experimental procedures were approved by the French Ministry of Education and Research, after consultation with the Ethical Committee #59 (authorization number: APAFIS#25932-2020060813556163 v7).

### Surgery

Adult mice (at least 8 weeks of age) were anesthetized using isoflurane anesthesia (induction 3 to 4%, maintenance 1 to 1.5%). Suppression of paw withdrawal, whisker movement, and eye-blink reflexes was used to control for the depth of anesthesia. Mice received anti-inflammatory medication (Meloxicam (8mg/kg), subcutaneously), were placed on a heating blanket with the head stabilized by a custom-designed nose clamp. Their eyes were kept moist with Ocry-gel (TVM Lab). Following a local subcutaneous injection of lidocaine (4 mg/kg), the skin covering the skull was cut, and Povidone-iodine 5% was applied to treat the surgical wound. A drop of 3% hydrogen peroxide was applied to the skull surface, and the skull was scraped with a scalpel. An oval-shaped craniotomy with a 6 mm longer axis was performed above the left somatosensory and motor cortices. A 6 mm diameter transparent glass window was glued in place using gel cyanoacrylate glue. A head fixation post was glued on the occipital bone. Dental cement was used to cover the skull and secure the head post and cranial window. Mice were monitored daily post-surgery until their weight returned to the pre-surgery level.

### Calcium imaging and real-time wave tracking

An imaging and real-time image-processing setup (Figure 1A) was assembled (R&D Vision, France) from a JAI GO-2400M-PMCL camera mounted on a custom-built tandem-lens epifluorescence macroscope (Ratzlaff and Grinvald, 1991) that combined two photographic lenses in a face-to-face configuration: a 50 mm f/1.2 Nikkor lense (sample side), and a 12.5 – 75 mm f/1.2 Fujinon zoom lens (camera side). Recordings were performed at 200 Hz with two alternating light sources: green, mounted on the side of the optical path (Thorlabs M530L4 green LED light source with a Thorlabs SM1U25-A collimator and Thorlabs LEDD1B driver) and blue, mounted on the main optical path (SciMedia Ltd. LEX3-Blue LED light source with a Chroma ET480/40x filter). Separation of the blue source and camera light paths was achieved with a dichroic mirror (Semrock FF495-Di03 reflecting all wavelengths < 495 nm). A band-pass filter (Edmund Optics 525/45, OD6) restricted to green the spectrum of light that reached the camera. The blue light excitation was used for GCaMP6f fluorescence excitation, while the green light source aimed to capture hemodynamic-related signals that contaminate the GCaMP6f fluorescence. To correct the GCaMP6f signal from these intrinsic contaminants, each image acquired under blue illumination was divided in real-time by the subsequent image taken under green illumination, generating a new sequence of frames at 100 Hz. To prevent high round-off error during division of integer-valued images, each blue image was multiplied by 64 before division.

After the correction for hemodynamics, fluorescence variation (dF/F_0_) images were obtained relative to a baseline fluorescence image F_0_, calculated from the 100 frames preceding the current frame. A 2D Mexican wavelet was applied to the data in the spatial dimension to retain components of cortical activity with sizes between 100 µm and 1 mm. After wavelet application, Otsu’s method (Otsu 1979) was used to create a binary mask corresponding to blobs of activity, and the local maximum within each blob, representing either a stationary or moving wave, was identified.

For each frame, all local maxima detected in that frame were compared with the last known coordinates of all tracked waves during the preceding ten frames. A local maximum detected in the current frame was considered part of an existing tracked Wave if the distance between the current local maximum and the last known coordinate of the Wave (within the preceding ten frames) was minimal compared to other Waves and not greater than ten pixels (0.2 mm), and if the difference in relative fluorescence value was less than an arbitrary chosen threshold. If no such match was found, the detected maximum was considered a new Wave to be tracked, as soon it was detected in at least three frames. Conversely, if no new match was found for a previously tracked Wave for ten frames, that Wave was considered to have ended.

To be considered a Conditioned Wave (CW), a Wave must have been detected first in the Start zone, in at least one frame, and then be tracked until it is detected in the End zone for at least two frames. Each new Conditioned Wave opened immediately a 500 ms opportunity window for reward delivery. Each subsequent frame in which the wave continued to meet the reinforcement condition extended the opportunity window by 10 ms. All licks detected inside the opportunity window were rewarded. If at least one lick was detected, the CW was labeled as Rewarded. The latency between image acquisition and opportunity window opening in the case of Conditioned Wave detection was less than 10 ms. The system was able to track up to 10 Waves simultaneously, and multiple Waves could be detected as Conditioned Waves simultaneously. Their opportunity windows would then overlap.

### Cortical mapping

After the mouse fully recovered from surgery (minimum of 3 weeks), the right hindlimb and forelimb were stimulated with a rod actuated by a piezoelectric bender, to localize the hindlimb- and forelimb-related regions in the somatosensory cortices of the anesthetized mouse (isoflurane 0.75–1.5%). The widefield calcium signal from 45 stimulations was recorded and averaged. The frame acquired 350 ms after stimulus onset was filtered using a 30x30 2D median filter. An extended-maxima transform (h = 0.01) was then applied to segment and identify the contour of the zone of evoked response.

### Habituation protocol and setting of initial task parameters

Once the mouse was trained to stay head-fixed in the setup, it was placed on a water restriction protocol and trained to receive water from a lick-port. Spontaneous activity was collected to determine the placement of the Start and End zones so that 1-5 Conditioned Waves spontaneously occurred per minute. For 15 of the 16 mice, there was no gap between the Start and End zones in the initial training protocol. For mouse #13, a 130 µm gap was present between the zones since the start of the training.

### Training Protocol

Water-regulated mice were placed in the setup, and an image of the brain was taken under blue illumination. Alignment of the Start and End zones between the current and previous sessions was performed using phase correlation registration. The training protocol continued for 10 – 30 minutes per session, with one session per day. During the training session, water droplets were delivered by a lick-port upon lick detection by a capacitive sensor within the reward opportunity window. During the session, we recorded the times of licks, opportunity window opening times (t_CW_), reward delivery times, real-time detected Wave trajectories, calcium signals and body movement (see below) synchronously. Only the first 10 minutes of session time were used for analysis, when the highest motivation was apparent.

### Analysis of behavioral performance

In order to identify the specific sessions during which a mouse performance was significantly better than baseline, we computed for each session the distribution of the time intervals between Rewarded Conditioned Waves (Figure 1G). A reference, *chance performance* distribution was defined by cumulating the intervals between Rewarded CWs observed during training sessions 1 and 2. A Kruskal-Wallis test was then applied to the set of distributions of intervals between Rewarded CWs computed across training sessions. If this first test identified that not all distributions of intervals between Rewarded CWs had the same distribution (p < 0.05), we then applied a Tukey-Kramer multiple comparison test to identify the sessions with a distribution significantly shorter (i.e., Rewarded CWs appeared more frequently) than that of sessions 1 & 2 (p < 0.05). Only time intervals longer than 500 ms were considered, corresponding to the duration of the reward opportunity window.

### Recording and quantification of body movement

During the training and habituation sessions, body movement was recorded at 100 Hz using a DMK33UX273 camera with a Fujinon DF6HA-1B 1:1.2/6 mm lens, equipped with an EFFI-RLSW-00-050-IR infrared ring light. The camera was synchronized with the blue light source used for calcium imaging. The right-side (contralateral to the craniotomy) hindlimb and forelimb were tracked using DeepLabCut (version 2.1.8.2) employing a ResNet-101 convolutional neural network (Mathis et al. 2018). Two points were tracked per limb, at its proximal and distal aspects. For each frame, the x,y coordinates of each limb were calculated as the average of the corresponding proximal and distal points. The pixel coordinates were converted into millimeters based on the headpost length located in the median plane, used as a reference. To reduce potential detection noise, a Kalman filter (acceleration variance 9×10⁶ mm²/s⁴) was applied to each limb coordinate to estimate position and instantaneous velocity. A generic instantaneous "Limb speed" was computed as the average of the absolute speed of the hindlimb and forelimb. Velocity peaks above 15 mm/s were detected to identify individual movements. For each detected movement, we searched from the peak, in both time directions, until velocity fell below 4 mm/s. Two lines passing through the peak and these points defined the movement onset and offset as their crossings with zero.

For mice #5 and #6, body movements were not recorded during the first training session; therefore, they were excluded from the initial movement analysis (Figure 4) but not from movement analysis after opening the gap between Start and End zones (Figure 5).

### Opening the gap between Start and End zones

Nine of the 10 mice attributed to the Learning group were involved in the Gap condition (Mouse #16 was not trained in this additional condition). The gap opening was performed after an average of 21 sessions on the initial protocol. For 7 of the 9 mice, the Start zone remained unchanged and the End zone was shifted along the task main axis (as defined in Figure 2I) by 130 µm to create the gap (Figure 5A), while for mice #5 and #6, the Start and End zones remained fixed and a 130 µm gap was added in between, thereby reducing their width. In addition, while for 7 of the 9 mice, the gap was introduced using the same task Main axis as in the No Gap condition, in two mice the location of the Start and End zones was modified before opening the gap: in mouse #11, the Start and End zones were inverted for 14 sessions, and in mouse #14, both zones were rotated around their common center by 32° for 10 sessions.

### Perfusion and Histology

The mice were anesthetized using isoflurane anesthesia (induction 3 to 4%, maintenance 1 to 1.5%) and placed on a stereotaxic frame. After removal of the implanted window, DiI fluorescent marker was injected at a depth of 1 mm at the corners of the Start and End zones. Following the administration of an overdose of pentobarbital (150 mg/kg), mice were perfused transcardially with saline followed by paraformaldehyde (4% in 0.1 M phosphate buffer). After overnight post-fixation of the brain in paraformaldehyde, 80 µm-thick coronal brain slices were exposed to 1:1000 DAPI for 5 minutes.

## ACKNOWLEDGMENTS

Experimental assistance and technical expertise was provided by Guillaume Hucher. We thank Thomas Deneux (NeuroPSI), Valentine Dhers and Olivier Lambert (R&D-Vision, France) for hardware and software development for the behavioral setup; Rodrigo Cofre and Alain Destexhe for their feedback and discussions throughout the project; Clément Picard, Rumeysa Can, and Konstantina Xanthopoulou Koukogia for their contribution to data analysis.

This work was funded by CNRS (CNRS 80|Prime, PRIME interdisciplinary label), Fondation pour la Recherche Médicale, La Fondation Dassault Systèmes, Agence Nationale pour la Recherche (ANR JCJC Mesobrain, ANR PRC Expect, Motorsense, Hermin, PROFouNd, PerBaCo, ANR PRCI ProPerO), Lidex NeuroSaclay, Idex Brainscopes, iCODE, hCODE.

