## Supplementary Movie 1 caption for "Goal-directed shaping of cortical waves"

**Supplementary Movie 1. Example cortical wave.**
Average raw relative GCaMP6f signal after hemodynamics artefact correction across the cortical window in mouse #14, aligned on $t_{CW}$, during the best performance session (n = 43 CWs).
